## Supplementary Figures S1 to S8 for "An Integrated Stress Response-independent role of GCN2 prevents excessive ribosome biogenesis and mRNA translation"

### SUPPLEMENTARY INFORMATION
### for

#### An Integrated Stress Response-independent role of GCN2 prevents excessive ribosome biogenesis and mRNA translation

**Authors**
Mónica Román-Trufero[1,2,3], Istvan T. Kleijn[4], Kevin Blighe[5], Jinglin Zhou[2], Paula Saavedra-García[2], Abigail Gaffar[2], Marilena Christoforou[2], Axel Bellotti[1], Joel Abrahams[6], Abdelmadjid Atrih[7], Douglas Lamont[7], Marek Gierlinski[8], Pooja Jayaprakash[9], Audrey M. Michel[9], Eric Aboagye[6], Mariia Yuneva[3], Glenn R. Masson[10], Vahid Shahrezaei[4] and Holger W. Auner[1,2,3,11]*

**Affiliations**
[1]Division of Haematology and Central Haematology Laboratory, Lausanne University Hospital (CHUV), CH-1011 Lausanne, Switzerland.
[2]Hugh and Josseline Langmuir Centre for Myeloma Research, Department of Immunology and Inflammation, Imperial College London, W12 0NN London.
[3]The Francis Crick Institute, NW1 1AT London, UK.
[4]Department of Mathematics, Imperial College London, SW7 2AZ London, UK.
[5]Clinical Bioinformatics Research, W2 6HP London, UK.
[6]Department of Surgery and Cancer, Imperial College London, W12 0NN London, UK.
[7]FingerPrints Proteomics Facility, School of Life Sciences, University of Dundee, DD1 5EH Dundee, UK.
[8]Data Analysis Group, Division of Computational Biology, School of Life Sciences, University of Dundee, DD1 5EH Dundee, UK.
[9]EIRNA Bio Ltd, T12 Y337 Cork, Ireland.
[10]Division of Cellular & Systems Medicine, School of Medicine, University of Dundee, DD1 9SY Dundee, UK.
[11]Faculty of Biology and Medicine, University of Lausanne, CH-1011 Lausanne, Switzerland.

This file includes:
Figures S1 to S8

#### Figure S1

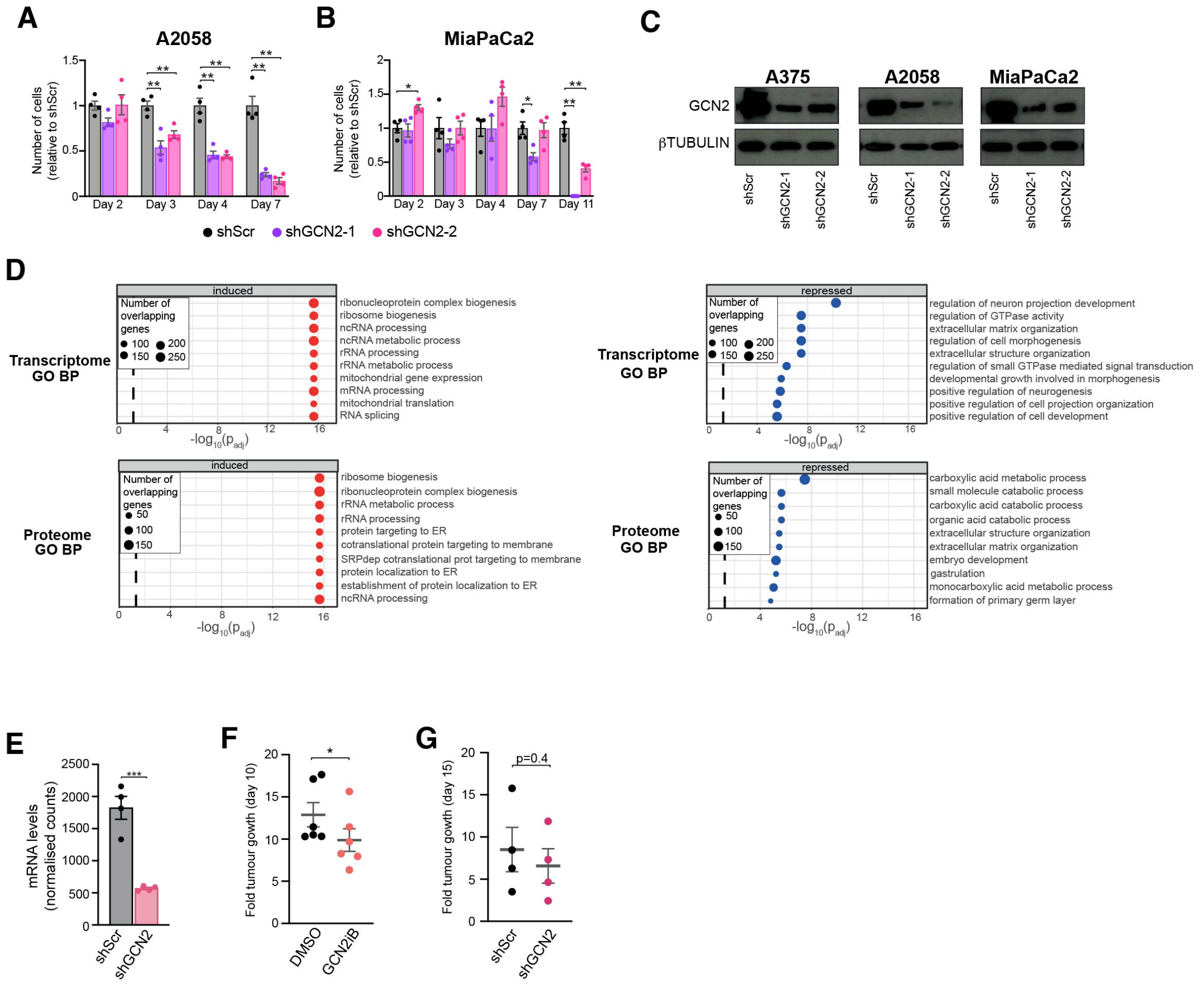


**Fig. S1. Transcriptome and proteome changes induced by GCN2 inhibition or depletion.** **(A)** Viability of A2058 cells after GCN2 knock down. Data shown as mean ± SEM of 4 independent experiments. Statistical significance was determined by two-way ANOVA and Dunnett's test for multiple comparisons. **(B)** Viability of MiaPaCa2 cells after GCN2 knock down. Data shown as mean SEM of 4 independent experiments. Statistical significance was determined by two-way ANOVA and Dunnett's test for multiple comparisons. **(C)** Immunoblot analysis of GCN2 levels in A375, A2058 and MiaPaCa2 cells for days after transduction with lentiviral particles carrying one of two different shRNAs against GCN2 or scramble control (Scr). **(D)** Top 10 induced and repressed Gene Ontology (GO) Biological Process (BP) terms in the transcriptome and proteome of A375 cells after treatment with GCN2iB (48h, 1 μM). **(E)** GCN2 transcript levels as determined by RNA-seq in xenografted tumours generated with A375 cells carrying a shRNA against GCN2 or a scramble control. Data shown as mean ± SEM of 4 mice. **(F)** Fold tumour growth relative to tumour volume at baseline after 10 days of treatment with 10mg/kg GCN2iB. Data shown as mean ± SEM of 6 mice. **(G)** Fold tumour growth relative to tumour volume at baseline after 15 days of shRNA induction with doxycycline. Data shown as mean ± SEM of 4 mice. (E) to (G) statistical significance was determined by unpaired t-test. *p<0.05, **p<0.005, ***p<0.001, ****p<0.0001.

#### Figure S2

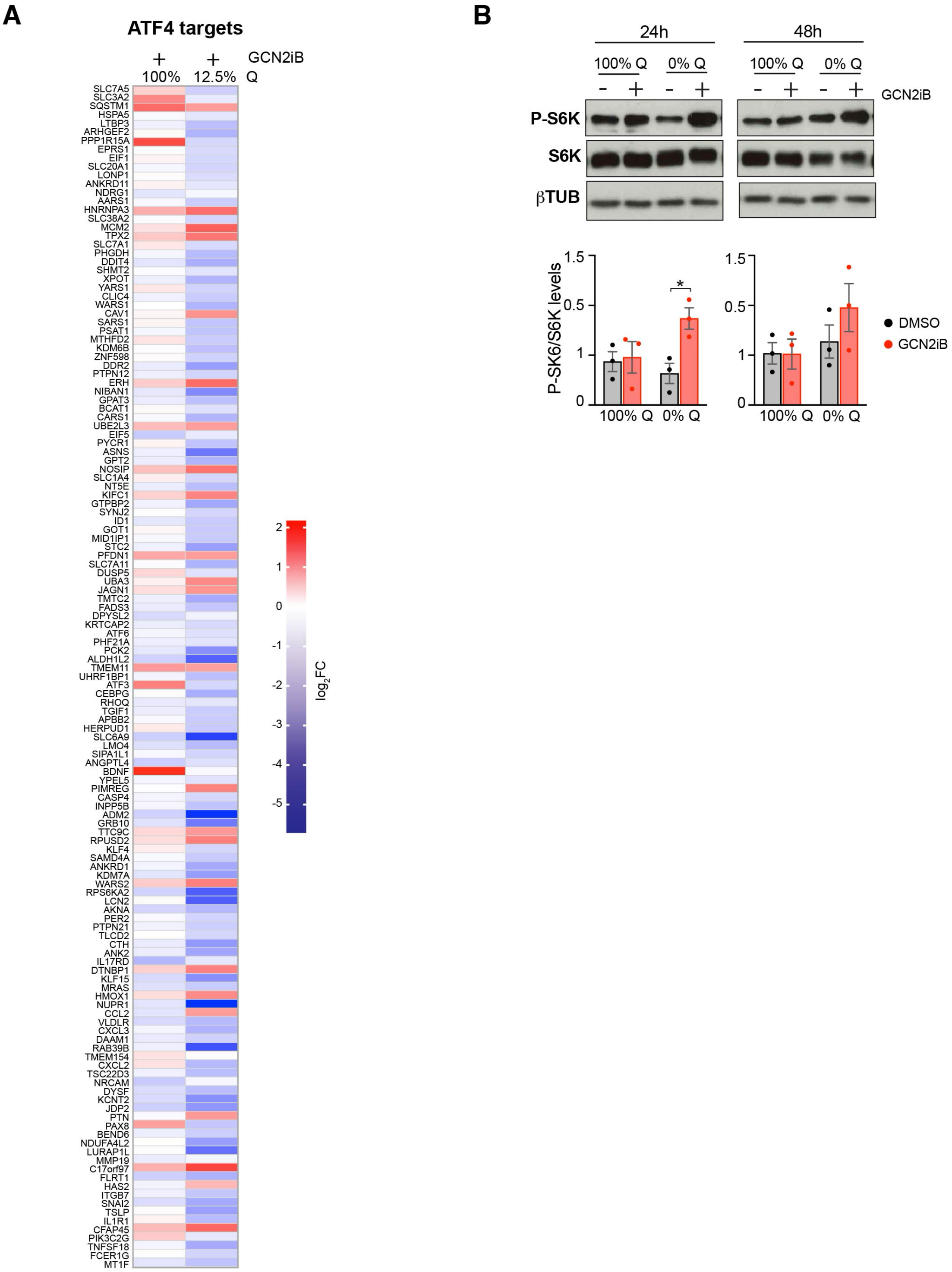


**Fig. S2 option B. Transcriptomic response to GCN2 inhibition depends on nutrient context.**

**(A)** Heatmap representation of ATF4 targets transcript level changes as determined by RNA-seq. Data are shown as $log_2$ fold change relative to DMSO control, n=4. A375 cells were cultured in complete medium (2 mM glutamine, 100%Q) or in glutamine-depleted medium (250 nM glutamine, 12.5%Q) and in the presence of 1 μM GCN2iB or vehicle control (DMSO). Samples were collected after 48h and processed for RNA sequencing and immunobloting. **(B)** Immunoblot analysis of S6 kinase phosphorylation in A375 cells grown in complete medium

(2 mM glutamine, 100%Q) or glutamine-depleted medium (0%Q) and treated for 24h and 48h with 1 μM GCN2iB (top), and bar graphs showing quantification of phosphorylated S6K relative to total S6K. Data shown as mean ± SEM of 3 independent experiments. Statistical significance was determined by two-way ANOVA and Sidák's test for multiple comparisons.

**Figure S3**

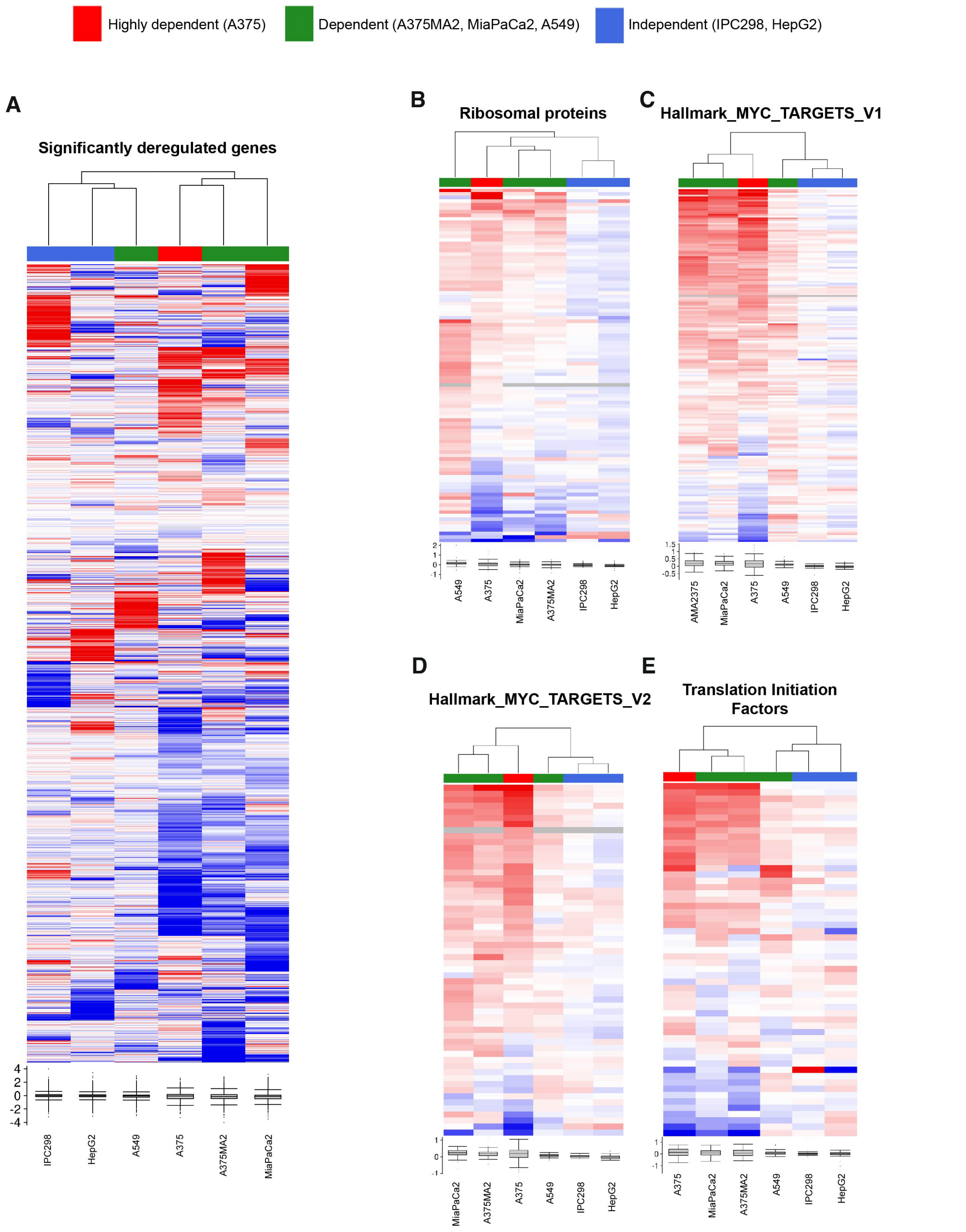


**Fig. S3. Cluster analysis identifies differences in response to GCN2 inhibition in GCN2-dependent and independent cell lines.**

Heatmap visualisations of hierarchical clustering analysis of RNA-seq data obtained from 6 cancer cell lines (IPC298, HepG2, A549, A375MA2, MiaPaCa2 and A375) of different tissue origin and with different degree of GCN2 dependency. Clustering performed based on transcript level changes upon 48h GCN2iB treatment on all significantly deregulated genes

**(A)**, ribosomal proteins **(B)**, the Hallmark categories MYC_targets_V1 **(C)** and MYC_targets_V2 **(D)**, and translation initiation factors **(E)**.

#### Figure S4

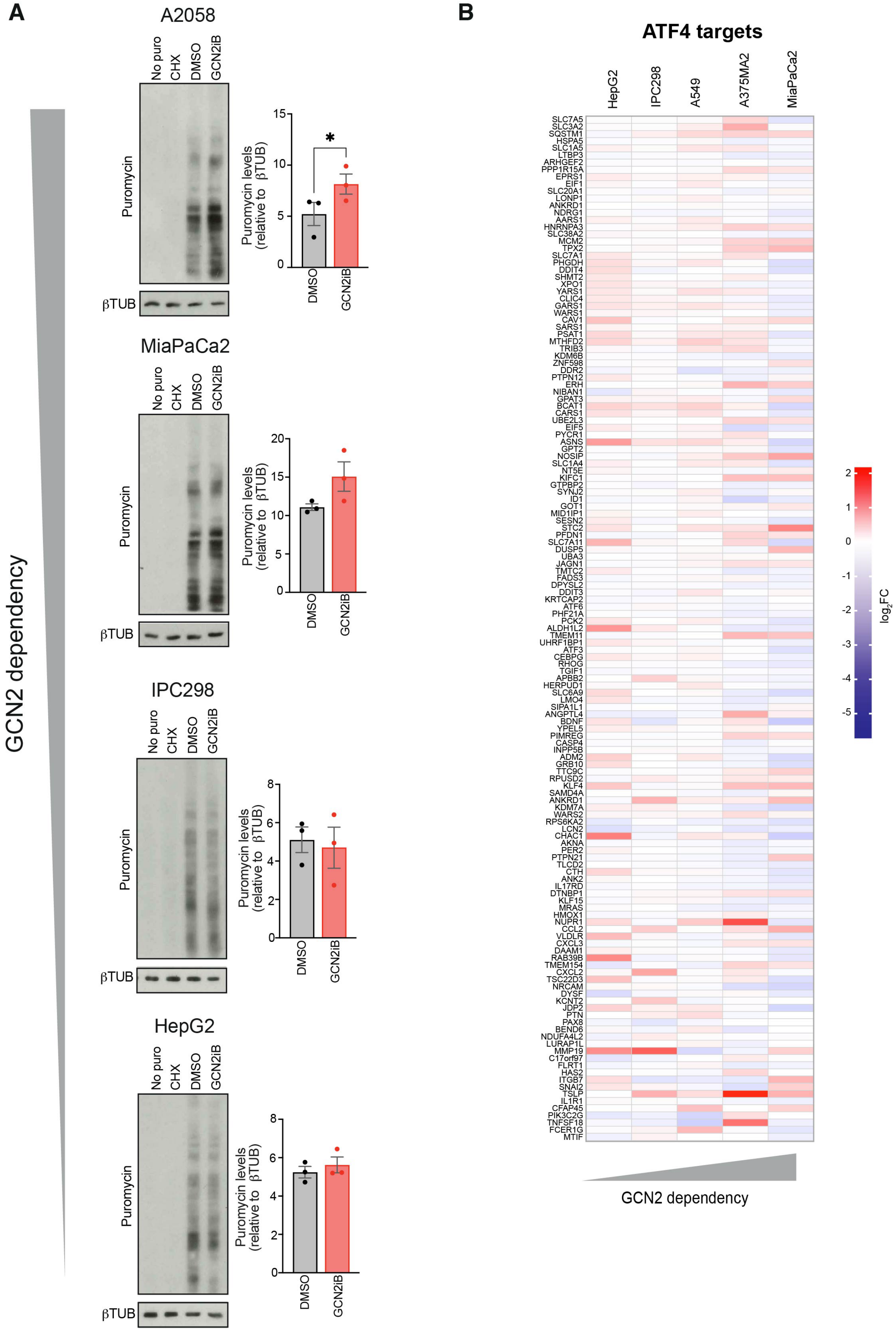

A
A2058
No puro
CHX
DMSO
GCN2iB
Puromycin
βTUB
Puromycin levels (relative to βTUB)
*
DMSO
GCN2iB
MiaPaCa2
IPC298
HepG2
GCN2 dependency
B
ATF4 targets
HepG2
IPC298
A549
A375MA2
MiaPaCa2
SLC7A5
SLC3A2
SQSTM1
HSPA5
SLC1A5
LTBP3
ARHGEF2
PPP1R15A
EPRS1
EIF1
SLC20A1
LONP1
ANKRD1
NDRG1
AARS1
HNRNPA3
SLC38A2
MCM2
TPX2
SLC7A1
PHGDH
DDIT4
SHMT2
XPO1
YARS1
CLIC4
GARS1
WARS1
CAV1
SARS1
PSAT1
MTHFD2
TRIB3
KDM6B
ZNF598
DDR2
PTPN12
ERH
NIBAN1
GPAT3
BCAT1
CARS1
UBE2L3
EIF5
PYCR1
ASNS
GPT2
NOSIP
SLC1A4
NT5E
KIFC1
GTPBP2
SYNJ2
ID1
GOT1
MID1IP1
SESN2
STC2
PFDN1
SLC7A11
DUSP5
UBA3
JAGN1
TMTC2
FADS3
DPYSL2
DDIT3
KRTCAP2
ATF6
PHF21A
PCK2
ALDH1L2
TMEM11
UHRF1BP1
ATF3
CEBPG
RHOG
TGIF1
APBB2
HERPUD1
SLC6A9
LMO4
SIPA1L1
ANGPTL4
BDNF
YPEL5
PIMREG
CASP4
INPP5B
ADM2
GRB10
TTC9C
RPUSD2
KLF4
SAMD4A
ANKRD1
KDM7A
WARS2
RPS6KA2
LCN2
CHAC1
AKNA
PER2
PTPN21
TLCD2
CTH
ANK2
IL17RD
DTNBP1
KLF15
MRAS
HMOX1
NUPR1
CCL2
VLDLR
CXCL3
DAAM1
RAB39B
TMEM154
CXCL2
TSC22D3
NRCAM
DYSF
KCNT2
JDP2
PTN
PAX8
BEND6
NDUFA4L2
LURAP1L
MMP19
C17orf97
FLRT1
HAS2
ITGB7
SNAI2
TSLP
IL1R1
CFAP45
PIK3C2G
TNFSF18
FCER1G
MTIF
2
1
0
-1
-2
-3
-4
-5
log2FC
GCN2 dependency

**Fig. S4. Differential regulation of protein synthesis upon GCN2 inhibition in GCN2-dependent and independent cell lines.**

**(A)** Immunoblot analysis of puromycinylated proteins in the indicated cell lines (left), and bar graphs showing quantification of puromycinylated proteins (right). Data shown as mean ± SEM of 3 independent experiments. Statistical significance was determined by paired t-test. **(B)** Heatmap representation of ATF4 targets transcript level changes as determined by RNAseq of the indicated 6 cell lines. Data are shown as $\log_2$ fold change, n=4. In (A) and (B), cells were treated with 1 μM GCN2iB for 48h. *$p<0.05$.

### Figure S5

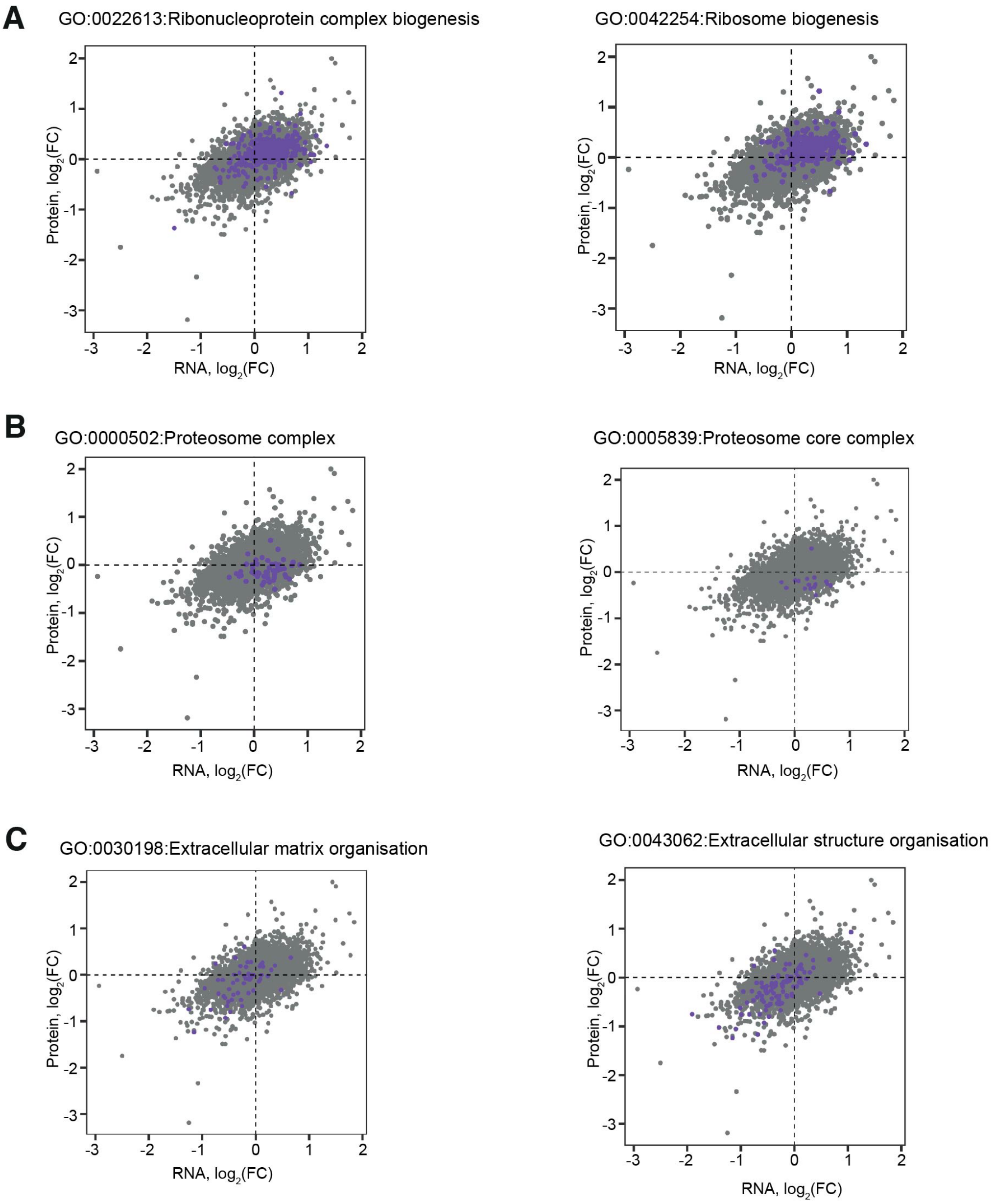


**Fig. S5. Differential regulation of functional gene/protein groups.**

Correlation plots showing the $log_2FC$ for each mRNA (X-axis) and protein (Y-axis) in A375 cells treated with 1 μM GCN2iB for 48h, genes belonging to Gene Ontology (GO) categories of interest are coloured in purple: **(A)** GO:0022613 Ribonucleoprotein complex biogenesis (left) and GO:0042254 Ribosome biogenesis (right); **(B)** GO:0000502 Proteasome complex (left) and GO:0005839 Proteasome core complex (right); **(C)** GO:0030198 Extracellular matrix organization (left) and GO:0043062 Extracellular structure organization (right).

#### Figure S6

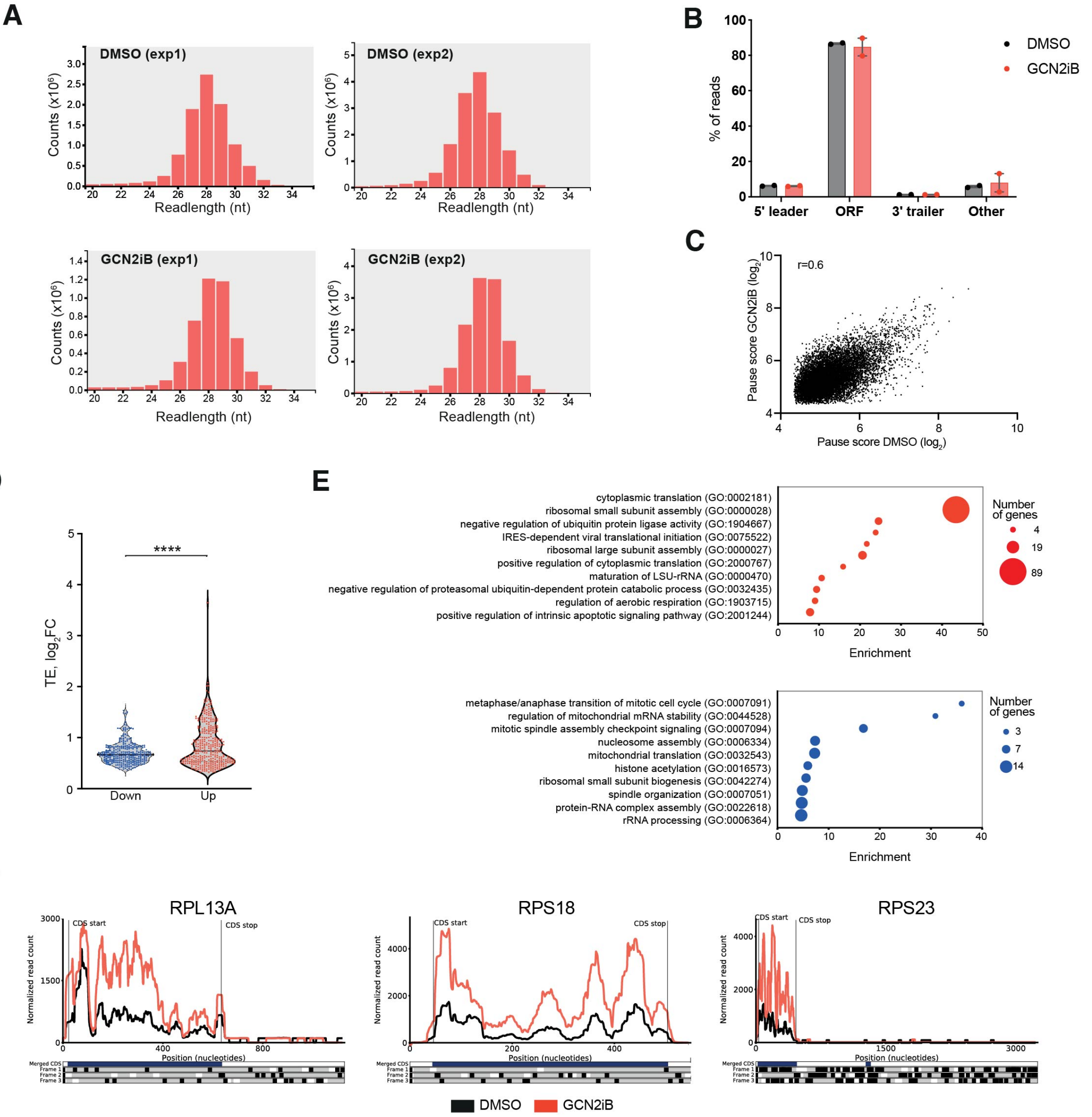


**Fig. S6. Ribo-seq quality control and ribosomal protein translation**

**(A)** Plots showing the expected read length distribution of the ribosomal footprints for the 4 ribo-seq experiments performed in this study. **(B)** Chart showing the percentage of reads aligning to the 5'UTR, the ORF, the 3'UTR or non-coding regions (other). **(C)** Comparison plot of the mean pause score of 10095 individual sites for A375 cells treated with GCN2iB (Y axis) vs vehicle control (X axis). **(D)** Violin plot showing the $log_2FC$ in translation efficiency (TE) in A375 cells treated with 1 μM GCN2iB for 48h, the black line indicates the median. **(E)** Bubble plots depicting the top 10 enriched Biological Process (BP) Gene Ontology (GO) terms of genes with increased (top) and reduced (bottom) TE in GCN2iB-treated A375 cells (1 μM 48h). **(F)** Comparison profiles showing ribosome occupancy along three representative ribosomal mRNAs in A375 cells treated with GCN2iB (red) or DMSO (black). ****p<0.0001.

#### Figure S7

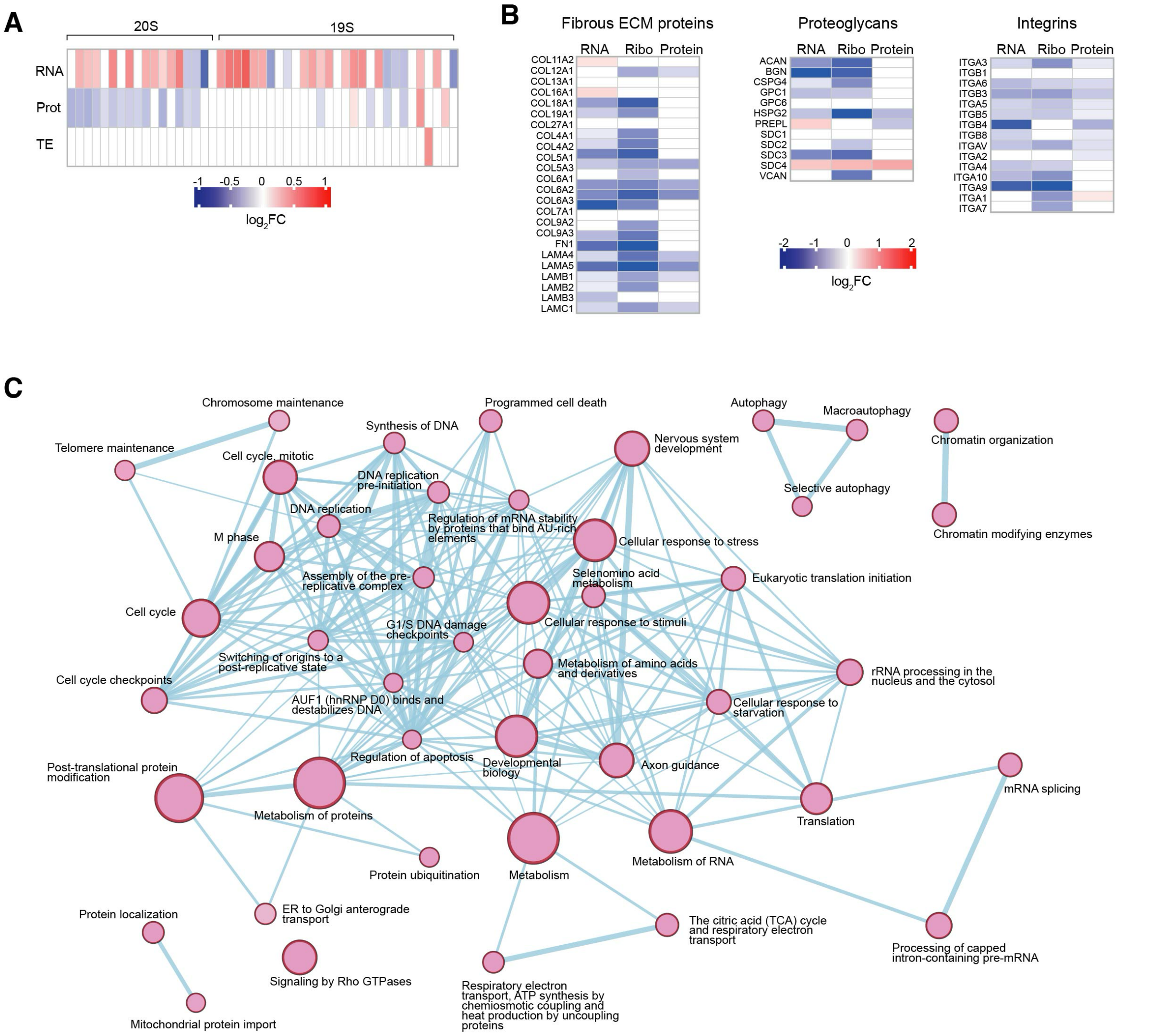


**Fig. S7. Regulation of translation by GCN2.**

**(A)** Heatmap representation of the changes in mRNA, protein abundance and TE for 20S and 19S proteasome subunit genes in GCN2iB-treated A375 cells (1µM 48h). **(B)** Heatmap representation of the changes in mRNA expression, ribosome density and protein abundance of extracellular matrix proteins and integrins in GCN2iB-treated A375 cells (1µM 48h). In A and B, data shown as mean of n=4 for mRNA, n=3 for protein and n=2 for ribosome density. **(C)** Network analysis of selected enriched Reactome pathways in GCN2iB-treated cells. Node size indicates the number of genes in the pathway and thickness of the edges the degree of overlap between pathways.

### Figure S8

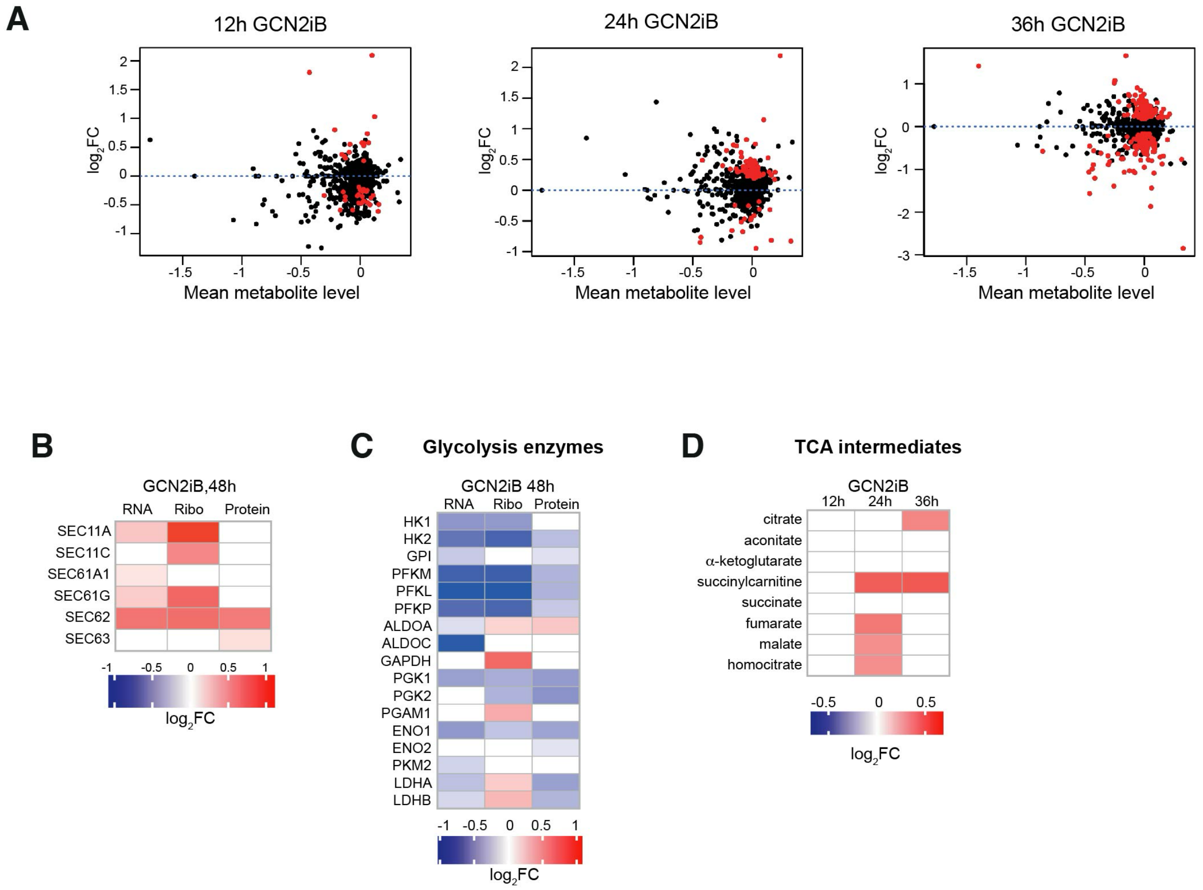


**Fig. S8. Effect of GCN2 inhibition on the metabolome of A375 cells.**
**(A)** Volcano plots showing the deregulated metabolites in A375 cells after GCN2 inhibition. Significantly (adjusted p value <0.05) deregulated metabolites are shown in red. **(B)** Heatmap representation of the changes in mRNA expression, ribosome density and protein abundance of subunits of the translocon and signal peptidase complexes in the ER. **(C)** Heatmap representations of the changes in mRNA expression, ribosome density and protein abundance of enzymes in the glycolysis pathway (B) and (C) Data shown as mean of n=4 for RNAseq, n=3 for proteomics and n=2 for Riboseq. **(D)** Heatmap showing the relative concentration of tricarboxylic acid cycle (TCA) intermediates in GCN2iB-treated A375 cells. Data shown as mean intensity normalised to DMSO, n=4. All panels show results in A375 cells treated with 1 μM GCN2iB for the indicated times.
